# Paying Attention to Speech: The Role of Cognitive Capacity and Acquired Experience

**DOI:** 10.1101/655274

**Authors:** Bar Lambez, Galit Agmon, Paz Har-Shai, Yuri Rassovsky, Elana Zion Golumbic

## Abstract

Managing attention in multi-speaker environments is a challenging feat that is critical for human performance. However, why some people are better than others in allocating attention appropriately, remains highly unknown. Here we investigated the contribution of two factors – Cognitive Capacity and Acquired Experience – to performance on two different types of Attention task: Selective Attention to one speaker and Distributed Attention among multiple concurrent speakers. We compared performance across three groups: Individuals with low (n=20) and high cognitive capacity (n=26), and Aircraft Pilots (n=25), who have gained extensive experience on both Selective and Distributed attention to speech through their training and profession. Results indicate that both types of Attention benefit from higher Cognitive Capacity, suggesting reliance on common capacity-limited resources. However, only Selective Attention was further improved in the Pilots, pointing to its flexible and trainable nature, whereas Distributed Attention seems to suffer from more fixed and hard-wired processing-bottlenecks.

## Introduction

Natural environments are characterized by an abundance of sounds bombarding the auditory system and competing for our attention. Focusing attention appropriately on relevant speech in adverse listening settings is a feat that many individuals find extremely challenging. Moreover, different contexts may require applying different attentional strategies. Some tasks require Selective Attention, i.e., focusing on a single speech-source while ignoring all others and avoiding distraction (Beaman, Bridges, & Scott, 2007; Broadbent, 1954; Cherry, 1953; Elliott & Briganti, 2012; Zion Golumbic et al., 2013), whereas other contexts require precisely the opposite: Distributing Attention among several speakers and gleaning semantic information from all of them (Baldock, Kapadia, & van Steenbrugge, 2018; Brungart, Kordik, & Simpson, 2005; Getzmann, Golob, & Wascher, 2016; Gygi & Shafiro, 2012). Although the term “Attention” is used to describe both processes, these tasks require overcoming fundamentally different perceptual and cognitive challenges. The primary challenge of Selective Attention is separating among acoustic sources and suppressing irrelevant portions of the auditory scene, a feat that becomes progressively difficult as the acoustic and linguistic overlap between concurrent speech increases (Bronkhorst, 2015; Drullman & Bronkhorst, 2004; Kidd et al., 2016; Neely & LeCompte, 1999; Oswald, Tremblay, & Jones, 2000). In contrast, the challenge of Distributed Attention stems from the limited nature of internal processing resources, that arguably pose inherent ‘bottlenecks’ for processing concurrent speech, the precise nature of which is still heavily debated (Bronkhorst, 2015; Deutsch & Deutsch, 1963; Driver, 2001; Lachter, Forster, & Ruthruff, 2004; Lavie, Hirst, de Fockert, & Viding, 2004)

The current study seeks to understand whether Selective and Distributed attention to speech rely on common underlying cognitive mechanisms and to highlight the similarities and distinctions between these two aspects of human attention. Specifically, we test the contribution of two factors – Cognitive Capacity and Acquired Experience – to performance on these tasks, as a mean for examining the processes underlying Selective and Distributed attention to speech.

Cognitive capacity is positively associated with performance on a range of attentionally-demanding tasks, and has been attributed to the availability of more cognitive resources as well as more effective top-down control over the allocation of these resources (Sörqvist & Rönnberg, 2014; Tsuchida, Murohashi, Katayama, & Murohashi, 2012; Wiemers & Redick, 2018). However, whether Selective and Distributed rely on shared attentional-resources, as proposed by some (Fusser et al., 2011; Kahneman, 1973; Salmela, Moisala, & Alho, 2014), or are differently affected by the availability of cognitive resources (Elliott & Briganti, 2012), remains currently unknown. Besides the contribution of cognitive capacity to performance, another fundamental question is how amenable attention is to improvement through training, and if it is possible to master attention-as-a-skill. Investigating whether performance can be further improved with experience touches upon the ‘fixed’ or ‘flexible’ nature of the processing bottlenecks and limitations they impose on performance.

In order to test the relative contributions of cognitive capacity and acquired experience to Selective and Distributed Attention to speech, here we compared performance on increasingly demanding attentional tasks, in three experimental groups. The role of cognitive capacity was assessed by comparing individuals with low- and high cognitive capacity, operationalized by an estimate of general intellectual functioning and working memory capacity. The role of acquired experience was studied by comparing performance of aircraft pilots with matched high cognitive capacity controls. Aircraft pilots afford a unique opportunity for studying the effects of acquired experience on attention, as their intense professional training and daily job pose high demands for both Selective and Distributed Attention to speech (Gopher, 1982; Gygi & Shafiro, 2012). Studying performance in this highly trained population allows us probe the limits of the speech-processing and attention systems, testing the extent to which humans are able to improve their abilities as well as identifying hard-wired bottlenecks that cannot be overcome.

## Methods

### Participants

Participants in this study were males between the ages of 24-49 (median = 29), with normal hearing and vision, and no history of psychiatric or neurological disorders or ADHD. They were recruited from three groups: Pilots, High Cognitive Capacity group (HC), and Low Cognitive Capacity group (LC; see Table 1). The Cognitive Capacity of each participant was evaluated by testing for general intellectual functioning (g-factor Test of Nonverbal Intelligence 3^rd^ Edition; TONI-3) and Working Memory Capacity (WMC, Wechsler Adult Intelligence Scale-Fourth Edition). Since these two measures are largely correlated (Borella, Pezzuti, De Beni, & Cornoldi, 2019), rather than distinguishing among them, here we consider them jointly as reflecting Cognitive Capacity.

**Table 1.** Participants Demographic characteristics

|  | Pilots (n=24) |  | HC (n=25) |  | LC (n=20) |  |
| --- | --- | --- | --- | --- | --- | --- |
|  | *M* | *SD* | *M* | *SD* | *M* | *SD* |
| Age (Years) | 29.42 | 5.52 | 29.16 | 4.4 | 29.7 | 7.25 |
| g-factor | 126.54 | 10.99 | 126.5 | 9.66 | 93.4 | 10.42 |
| Working Memory | 13.75 | 2.78 | 13.42 | 2.06 | 9.5 | 2.26 |

The Pilots group consisted of 25 active, commercial pilots, recruited from commercial airline companies, between the ages of 25-45. All Pilots had continuous flying experience of at least 7 years, allowing for the development of expertise (Ericsson, Krampe, & Tesch-Römer, 1993). The HC group consisted of 26 participants, who were matched to the Pilots on age (p=1.0), g-factor (p=0.992), and WMC (p=1.0). The LC group consisted of 20 participants who were also matched to the Pilots on age (p=1.0), but had significantly lower g-factor and WMC than the HC and Pilot groups (both p<0.001). Two participants (1 Pilot, 1 HC) were rejected from data analysis due to technical errors. Demographic data was collected using a self-report sociodemographic questionnaire.

The study was preregistered prior to commencement of data collection through the Center for Open Science (COS; https://osf.io/tk6gc), describing the specific experimental design and power-analysis for a-priori estimation of the required group-size. The original preregistration focused only on the HC and Pilot groups and recruitment of the LC group was decided upon at a later stage. The study was approved by the Ethics Committee of Bar Ilan University, and signed informed consent was obtained from each participant prior to the experiment. Participants were reimbursed for their time and travel expenses.

### Stimuli

The stimuli were comprised of a list of short Hebrew words, including monosyllabic nouns (e.g., pitcher; “*Kad*”) and digits (e.g., seven; “*Shevah*”). The words were recorded by two male and two female speakers, rendering the speakers clearly distinguishable. Audio editing of the individual words and their combination into sequences were performed using Audacity (www.audacityteam.org) and Matlab (The MathWorks, Inc). The perceived loudness of each speaker was equated offline and verified using inter-rater reliability testing. Word lengths varied between 800 ms to 1100 ms, and they were concatenated into 55-second long sequences, separately per speaker. The inter-stimulus-interval (ISI) between words varied across speakers between 600-750ms (but was held constant within speaker), in order to minimize common onsets and offsets. Sequences by different speakers were then combined to create diotic multi-talker scenes, according to the different experimental conditions, as explained below.

### Experimental procedure

We designed two parallel word target-detection tasks, probing Selective and Distributed Attention to speech under multi-speaker conditions (Figure 1). The Selective Attention task, required participants to attend to one designated (“attended”) speaker and respond to the target-word uttered only by this speaker. In the Distributed Attention condition, participants were instructed to respond to the target word spoken by <u>*any*</u> of the concurrently presented streams. The two tasks were matched for low-level acoustics, and the level of difficulty was parametrically varied by increasing the number of competing speakers from two to four. The same target word (tree; *“Etz”*) was used in all conditions and occurred 8 or 9 times per trial. In the Selective Attention condition additional ‘catch-targets’ could occur equi-probably in the unattended sequences, which participants were accordingly supposed to ignore.

Selective and Distributed trials were presented in four separate blocks (two per condition), in counter-balanced order across subjects. Each block consisted of 12 trials where the Number of Speakers was pseudo-randomized between 2 to 4, as well as two trials where only one speaker was presented (this condition was not included in the statistical analysis). The experiment was conducted in a sound-attenuated booth, and speech stimuli were presented through headphones (Sennheiser, HD 280 Pro). The experiment was programmed using Psychopy software (Peirce et al., 2019), and responses were recorded using a response box (Cedrus RB 840). Response Accuracy and Reaction Times (RTs) were extracted for each target and used for statistical analysis.

**Figure 1.**
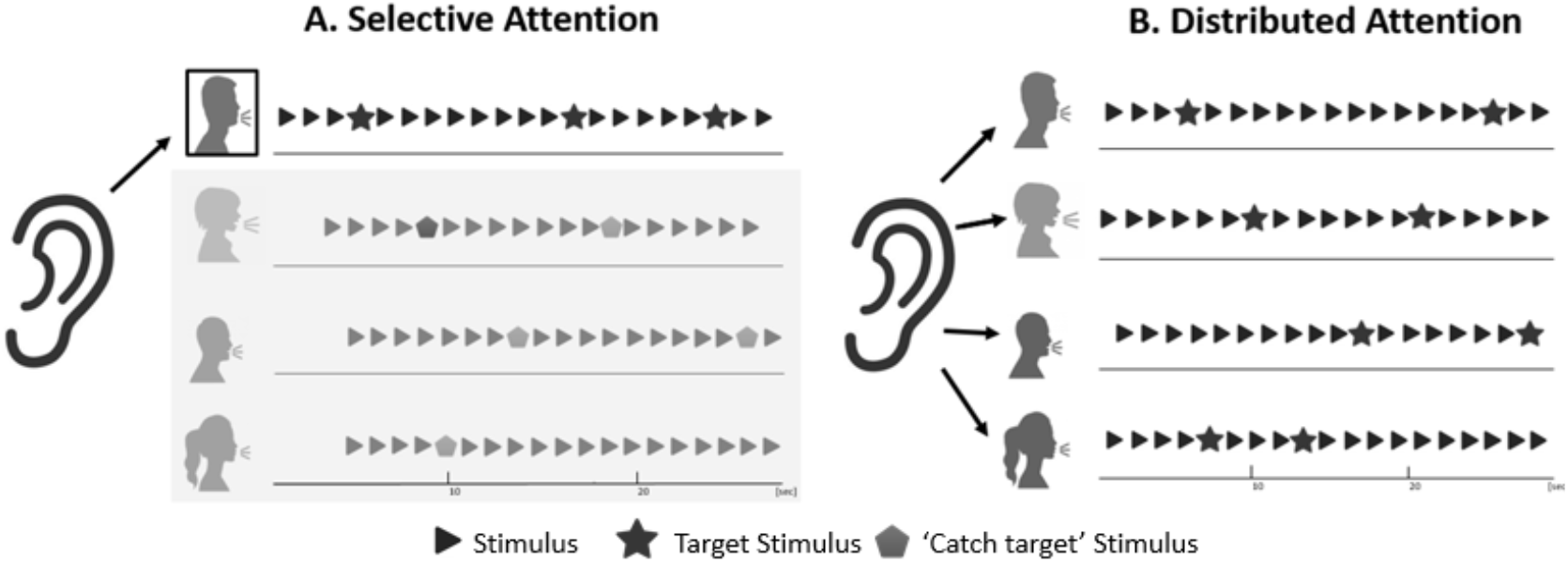
**A.** Example of the trial structure in the Selective Attention condition, in the case of four concurrent speakers. The speech stimulus of each speaker consisted of a sequence of Hebrew words (indicated by triangles). In Selective Attention blocks, participants were instructed to attend to one designated “attended” speaker and respond to a target-word (marked with star icon) uttered only by this speaker, while ignoring all the other speakers. A similar number of catch-targets (the target-word uttered by an unattended speaker, marked with a pentagon icon) were distributed among the other unattended speakers. The number of concurrent speakers varied between two and four, across trials. **B.** Example of the trial structure in the Distributed Attention condition, in the case of four concurrent speakers. The stimuli are similar to those used in A, however here participants were instructed to respond to the target-word uttered by *any* of speakers, requiring participants to distribute auditory attention among all concurrent speakers. Here too, the number of concurrent speakers varied between two and four, across trials.

#### Statistical Analysis

To test for differences between conditions, we fit the behavioral responses with mixed effects linear regression models, using R’s lme4 package (Bates, Maechler, Bolker, & Walker, 2015): for RT data we used a linear mixed effects regression and for the accuracy data we used a generalized linear mixed effects regression. The advantage of mixed effects models is that they account for variability between subjects and correlations within the data (Baayen, Davidson, & Bates, 2008); in the case of accuracy, they also allow accurate analyses of binomial data in repeated-measures designs (Jaeger, 2008).

The model fitted for the RTs included three factors and their interactions as fixed variables: Attention (*Selective, Distributed*), Group (*LC, HC, Pilots*) and Number of Speakers (*2, 3, 4*). The factor Attention was sum coded (i.e., the beta estimate represents deviance from the grand mean), the factor Group was treatment coded (i.e., one beta compares Pilots to HC and another beta compares LC to HC), and the factor Number of Speakers was difference coded (i.e., one beta codes for the difference between 3 and 2 speakers, and the second beta codes for the difference between 4 and 3 speakers). We included by-subject random intercepts, as well as by-subject random slopes for Attention and Number of Speakers. To better comply with the assumption of normality, we used the logarithmic transformation of RT as the dependent variable. As a pre-processing step, we removed from the data any RTs that were below 300 ms or above 2000 ms, thus removing 0.6% of the total data. P-values were obtained using the Satterthwaite approximation of degrees of freedom (Satterthwaite, 1946), which is implemented in R’s lmerTest package (Kuznetsova, Brockhoff, & Christensen, 2017). For accuracy, we used the same fixed variables and coding schemes. By-subject random intercepts and by-subject random slopes for attention were included. For hit trials in which RTs were either below 300 ms or above 2000 ms, accuracy was set to be 0. The reported p-values are based on asymptotic Wald tests which are included in the summary of R’s glmer function.

## Results

Performance was relatively high on all tasks, with an average of 90.5% detection rate across conditions and groups (±6.4% SD). Moreover, in the Selective Attention condition the false-alarm rate was extremely low (0.9%±1.3%; responses to the target-word spoken by unattended speakers), indicating that participants understood the difference between the Selective and Distributed tasks and followed instructions accordingly. Mixed-linear regression models were used to assess the effects of Attention Type (Selective vs. Distributed), Number of Speakers (2spk, 3spk and 4spk) and Group (LC, HC and Pilots) on accuracy and reaction times (RTs). Results across all conditions and groups are shown in Figure 2, and the full statistical results of all contrasts in the regression models are listed in Tables S1 & S2.

**Figure 2:**
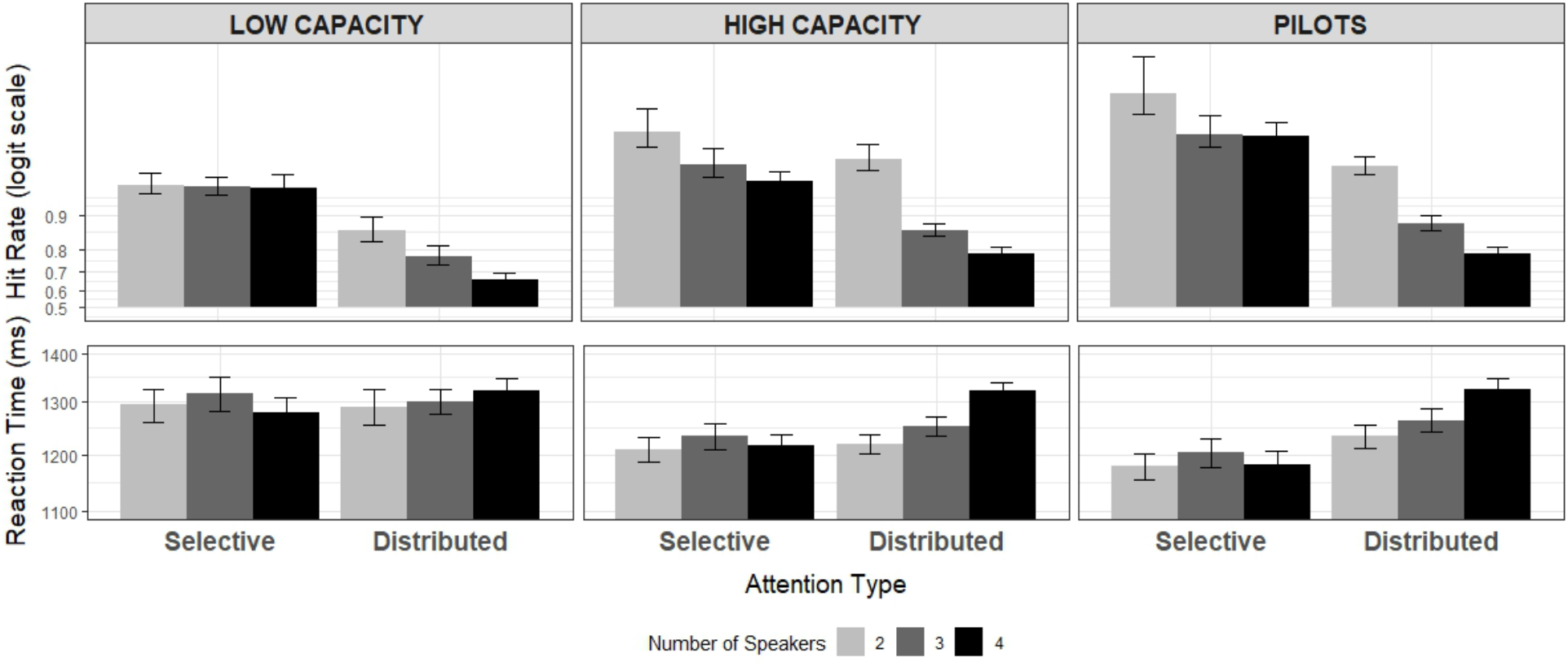
Mean Hit Rates (top) and RTs (bottom) on the Selective and Distributed Attention tasks, as a function of the number of concurrent speakers (2-4), for each of the three experimental groups: Low Cognitive Capacity, High Cognitive Capacity and airplane Pilots.

Comparison of performance between the two Attention Type tasks indicated that the Distributed Attention condition was more difficult than the Selective Attention condition, as manifest in reduced accuracy (β=-0.8, z=-5.6, p<0.0001) and prolonged RTs (β=0.02, t=3.5, p=0.0009). Performance also generally deteriorated as the number of concurrent speakers increased, observed in the 3spk vs. 2spk contrast (Accuracy: β=-1.3, z=-5.9, p<0.0001; RT: β=0.02, t=3.5, p=0.0008) and the 3spk vs. 4spk contrast (Accuracy: β=-0.5, z=-3.2, p=0.001; RT: β=0.02, t=3.3, p=0.001). Some of these effects interacted with the Attention Type, indicating that increasing the number of speakers adversely affected performance on the Distributed Attention task, and less so (or not at all) in the Selective Attention task [RTs (*4spk vs. 3spk) × Attention Type*: β=0.03, *t*=6.2, p<0.0001; Accuracy (*3spk vs. 2spk) × Attention Type*: β=-0.5, *z*=-2.1, p=0.03] (Figure 3).

**Figure 3:**
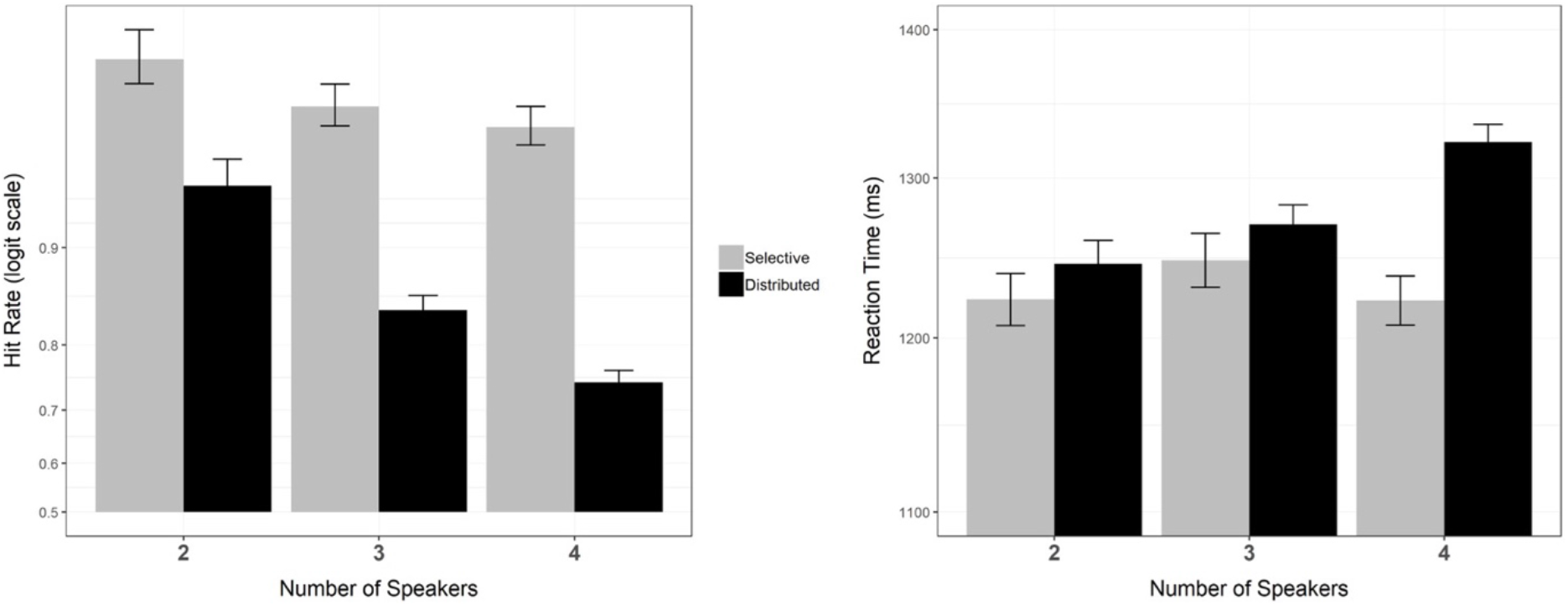
Mean Hit Rates (left) and RTs (right) on the Selective and Distributed Attention tasks, as a function of the number of concurrent speakers (2-4), collapsed across groups. Overall performance was worse in the Distributed Attention condition. Increasing the number of speakers had a more adverse effect on performance in the Distributed vs. Selective Attention condition. Error bars indicate SEM.

With regard to differences between the three Groups (Figure 4), the LC group demonstrated significantly lower accuracy than the HC group (β=-0.8, *z*=-3.9, p<0.0001), and a trend towards slower RTs (β=0.04, *t*=1.8, p=0.08). This contrast did not interact significantly with Attention Type, suggesting that both types of attention were similarly influenced by the difference in Cognitive Capacity between these two groups. Some additional higher-order interactions of the differences between the LC and HC groups with the Number of Speakers were also significant [(*LC vs. HC) × (2spk vs. 3spk)*: Accuracy β=0.9, *z*=3.5, p=0.0005; [(*LC vs. HC) × (3spk vs. 4spk)*: RTs: β=-0.02, t=-2.4, p=0.02]. These interactions likely stem from floor effects observed in the LC group, who displayed overall worse performance regardless of the number of speakers.

**Figure 4:**
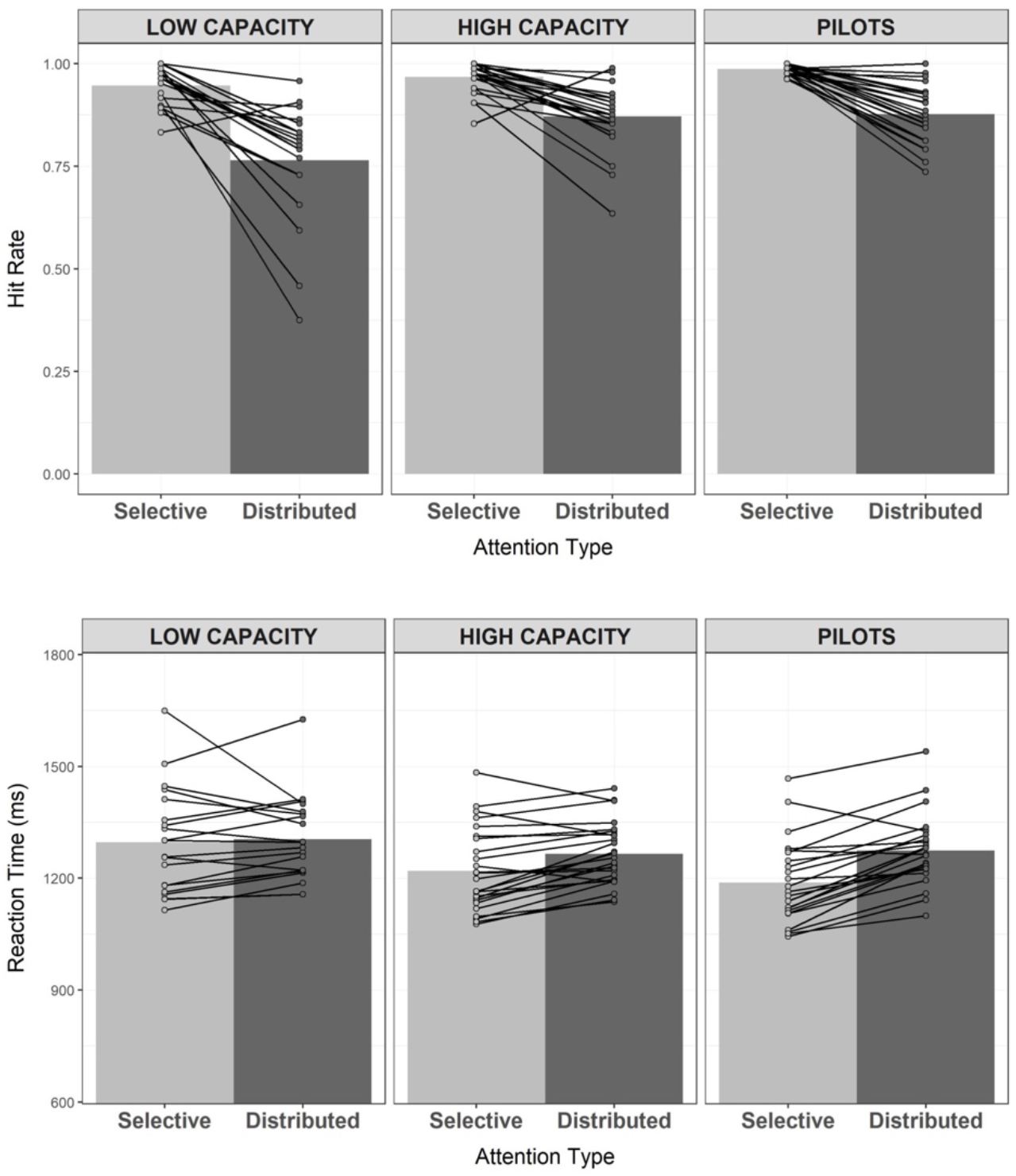
Mean accuracy (top) and RTs (bottom) in the Selective and Distributed conditions, separately for each of the three groups. Black lines connect between the scores of individual participants on the Selective and Distributed tasks.

Comparison of performance between the HC group and the Pilot group, did not reveal any main effects of Group on accuracy or RTs (RTs: p=0.7, Accuracy: p=0.07), however there was a significant interaction with Attention Type [*(HC vs. Pilots) × AttentionType*, RTs: β=0.02, *t*=2.4, p=0.02, Accuracy: β=-0.4, *z*=-2, p=0.05)]. This indicates that in the Selective Attention task the Pilot group performed better than their HC matched controls, however the two groups did not seem to differ in their performance on the Distributed Attention task.

## Discussion

The current study aimed at understanding the contributions of Cognitive Capacity and Acquired Experience to performance of Selective and Distributed Attention to speech. We find that both Selective and Distributed Attention are improved in individuals with high Cognitive Capacity, suggesting some reliance on common executive resources. However, these two facets of attention seem to differ in the processing bottlenecks encountered as well as in their fixed or flexible nature. Specifically, results suggest that Selective Attention is a highly effective and malleable process that can withstand substantial adverse conditions. In contrast, Distributed Attention suffers from more severe processing bottlenecks that are arguably of a fixed nature and are less prone to improvement through training.

The comparison between Selective and Distributed Attention indicates that not only is the latter more effortful (Baldock et al., 2018), but they are differently affected by increasing the number of concurrent speakers. For Selective Attention, we find only a slight decrease in target-detection accuracy and no impact on RTs as the number of concurrent speakers increased from two to four. This ability to withstand severe acoustic and informational masking, is in line with accounts suggesting that distractor-speakers are not fully segregated among themselves but rather treated collectively as acoustic ‘background’ (Hausfeld, Riecke, Valente, & Formisano, 2018; Puvvada & Simon, 2017). This pattern demonstrates a high degree of resilience of the auditory and/or attention system for dealing with background noise, as long as the cognitive task is limited to processing only one speaker. In contrast, Distributed Attention does not allow designating portions of the acoustic scene as ‘background’, and increasing the number of speakers also increases the perceptual and cognitive-load of the task itself (Baldock et al., 2018). This resulted in a sharper decline in accuracy and prolonged RTs as the range of Distributed Attention broadened from two to four speakers. This pattern indicates that attempting to process multiple concurrent speech-inputs draws upon perceptual and cognitive resources that are of a more limited nature, either at the level of the auditory system (Murphy, Spence, & Dalton, 2017) or at higher linguistic levels (Lachter et al., 2004).

The capability and limitations for parallel speech processing have been the focus of long-standing theoretical debates. However, much of this has been fueled by Selective Attention studies focusing on whether so-called unattended speech is processed (Beaman et al., 2007; Ding et al., 2018; Driver, 2001; Lachter et al., 2004; Röer, Körner, Buchner, & Bell, 2017; Wood & Cowan, 1995). The use of a Distributed Attention paradigm, as used here, provides a substantially more direct way for evaluating the ability and limitations of encoding, processing and responding to concurrent speech. Results highlight the fact that Distributed Attention suffers more substantially from processing ‘bottlenecks’ relative to Selective Attention, and provide insight into the nature of these bottlenecks. We now turn to discuss the role of Cognitive Capacity and Acquired Experience in mitigating these bottlenecks.

Both Selective and Distributed Attention require top-down control for allocating processing resources according to behavioral goals (Murphy et al., 2017). The Cognitive Capacity underlying top-down control varies across individuals and is strongly associated with a variety of executive processes, including selective, sustained and divided attention (Carroll, 1993; Kane & Engle, 2002). Previous studies have demonstrated that individuals with low WMC can be more susceptible to distraction (Hughes, 2014; Sörqvist & Rönnberg, 2014), incur a higher cost for attention switching (Lin & Carlile, 2015), display poorer speech-in-noise intelligibility (Keidser, Best, Freeston, & Boyce, 2015; Oberfeld & Klöckner-Nowotny, 2016), and exhibit overall slower processing speed (Fry & Hale, 1996). In line with these findings, we find that the LC group displayed poorer performance on both Selective and Distributed attention tasks, relative to the HC group. Moreover, there was no apparent difference in the effect of Cognitive Capacity on the two tasks. The plethora of attention-related deficits associated with reduced Cognitive Capacity is in line with perspectives suggesting that even though the behavioral goals and required cognitive operations may differ across attentional tasks, they nonetheless rely on overlapping attentional-resources and common neural substrates (Fusser et al., 2011; Humes, Lee, & Coughlin, 2006; Kahneman, 1973; Kiyonaga & Egner, 2013; Salmela et al., 2014; Salo, Salmela, Salmi, Numminen, & Alho, 2017),

Beyond an individual’s Cognitive Capacity, performance on difficult tasks can often improve substantially with training. Learning, and its accompanying neural plasticity, is one of the hallmarks of the nervous system, allowing organisms to adapt behavior to changing environmental and internal goals. Observing improvement by training is indicative of inherent flexibility of the underlying cognitive process (Buitenweg, Murre, & Ridderinkhof, 2012). Conversely, when no degree of training can improve performance beyond a certain level, this suggests a ‘hard bottleneck’, reflecting the upper limit of the system (Enns, Kealong, Tichon, & Visser, 2017). Comparison between the Pilot group and their matched HC controls provided a unique opportunity to assess whether the cognitive processes underlying Selective and Distributed Attention, are amenable to improvement through extensive training and experience. In the cockpit, Pilots need to meet extremely high standards of attention to multiple concurrent sources of speech, manifest both by monitoring several radio-channels (Distributed Attention), as well as focusing on one relevant radio-channel and disregarding irrelevant channels (Selective Attention). Their experience in performing these tasks builds on years of intense training and constant maintenance of these skills throughout their career (Hilburn, 2004). Indeed, it is difficult to imagine a more intense training regime for the refinement of attention skills to concurrent speech in humans, and consequently the Acquired Experience of Pilots substantially exceeds that gained through any experimental training program in both length and intensity.

Here, Pilots displayed an advantage on the Selective Attention task relative to their matched HC controls, which manifest in improved accuracy and faster RTs. This suggests that besides Cognitive Capacity, experience and training can improve Selective Attention to speech even further. In line with these results, previous studies have found improved Selective Attention performance in Pilots after successful completion of a two year long flight training program (Gopher, 1982), and similar advantages have been reported in air-traffic controllers (Arbula, Capizzi, Lombardo, & Vallesi, 2016). Critically, the amenability of Selective auditory attention performance to training is not limited to these highly-selective groups. Improvement in Selective Attention performance has been demonstrated after a 4-week dichotic-listening training program in healthy adults (Soveri et al., 2013), and similar training effects have been reported in typically developing children (Murphy, Moore, & Schochat, 2015), as well as children with dyslexia (Helland et al., 2018). These findings point to the flexible nature of auditory Selective Attention, which can be improved through formal training programs and as well as frequent practice (Mishra, de Villers-Sidani, Merzenich, & Gazzaley, 2014).

In contrast, our results suggest that Distributed Attention does not enjoy similar flexibility. Despite the rigorous training and Acquired Experience of Pilots, their performance was not significantly better their matched non-pilot controls. This points to an upper-limit on the ability to distribute attention among speech, that is not amenable to improvement through training. This pattern supports limited-resources perspectives of concurrent speech processing (Lachter et al., 2004) and suggests that bottlenecks may be hard-wired, possibly dictated by the a-priori availability of cognitive resources. It is important to acknowledge that the comparison between Pilots and the HC group performed here focused on individuals who can already perform the Distributed Attention task quite well, due to their high Cognitive Capacity. Future research should further look into possible effects of training on improving Distributed Attention performance in individuals with initially lower Cognitive Capacity.

To conclude, understanding why some people are better than others in allocating attention to speech in complex multi-speaker environments is extremely important, given its ecological significance to many real-life situations. This study elucidates the crucial role of cognitive capacity in the ability to process multiple concurrent speakers and distribute internal resources among them in accordance to task-goals. Moreover, it highlights the capacity and limitation of extensive training for improving different types of attentional performance. These results contribute to long-standing theoretical debates regarding the nature of ‘cognitive bottlenecks’ that limit the ability to deal with the influx of sensory information encountered in natural environments, and shed new light on their flexible or fixed nature.

## Acknowledgements

This work was supported by the following research grants: Marie Curie Career Integration Grant #631265 (EZG), and Binational Science Foundation Grant #2015385 (EZG).

## Author Contributions Statement

BL, YR & EZG designed the study, BL & PHS collected the data, BL & GA ran statistical analyses & prepared the figures, BL & EZG wrote the main manuscript text. All authors reviewed the manuscript.

## Data Availability

The data and code for statistical analysis in R are available on the Golumbic Lab website: www.golumbiclab.org/data

